# Lysosomal enzyme tripeptidyl peptidase 1 plays a role in degradation of beta amyloid fibrils

**DOI:** 10.1101/639682

**Authors:** Dana Cruz, Mukarram El-Banna, Amitabha Majumdar, David E. Sleat, Michelle Muldowney, Peter Lobel, Frederick R. Maxfield

## Abstract

Alzheimer’s disease (AD) is characterized by the accumulation of amyloid plaques surrounded by microglia. In cell culture, microglia internalize fibrillar β-amyloid but do not degrade it efficiently. Unactivated microglia have a relatively high lysosomal pH, which impairs the activity of lysosomal proteases. Previous studies showed that activation of microglia with macrophage colony stimulating factor decreases lysosomal pH and enhances fibrillar β-amyloid degradation. We investigated the role of the lysosomal protease tripeptidyl peptidase 1 (TPP1) in cell culture and in a mouse model of Alzheimer’s disease. Increased levels of TPP1 in unactivated microglia enhanced fibrillar β-amyloid degradation. Conversely, reduction of TPP1 led to decreased fibrillar β-amyloid degradation in activated microglia, macrophages, and other cells that degrade fibrillar β-amyloid efficiently. Reduction of TPP1 in an AD model mouse using a gene-targeted hypomorphic *Tpp1* allele increased plaque burden. These results suggest that decreased TPP1 potentiates AD pathogenesis and that strategies to increase TPP1 activity may have therapeutic value.

**Highlights:**

*In microglia, TPP1 is important for the degradation of fibrillar β-amyloid.
*Increased TPP1 in microglia results in enhanced fibrillar β-amyloid degradation.
*In an AD mouse model, reduction of TPP1 led to increased amyloid plaque deposition.

## 1. Introduction

Alzheimer’s disease (AD) is characterized by the accumulation of β-amyloid plaques and tau neurofibrillary tangles. In AD patient brains, β-amyloid plaques are surrounded by microglia, the resident immune cells of the central nervous system (Hopperton et al., 2018; Taipa et al., 2017). In culture, microglia take up small particles of fibrillar β-amyloid (fAβ) via Scavenger Receptor A (SRA) mediated endocytosis and deliver it to lysosomes but fail to degrade it efficiently (Chung et al., 1999; Majumdar et al., 2008; Masters et al., 2015; Paresce et al., 1996). In culture, untreated microglia continuously exposed to fAβ have lysosomes engorged with undigested fAβ (Paresce et al., 1997), resembling the deterioration of Aβ clearance with time at later stages of AD (ElAli and Rivest, 2016). Microglia do not lack the lysosomal enzymes important for macromolecule degradation (Chung et al., 1999; Majumdar et al., 2008; Paresce et al., 1997; Paresce et al., 1996). However, their average lysosomal pH of 6 is higher than that of macrophages and other cell types (Majumdar et al., 2007). The less acidic lysosomal pH of the microglia is suboptimal for many lysosomal proteases (Solé-Domènech et al., 2016).

Mass spectrometry analysis showed that the first twelve N-terminal amino acids (the unstructured free end) of fibrillar Aβ42 were degraded by microglia within 3 days, while degradation of the β-sheet region was very slow (Majumdar et al., 2008). In mice immunized against Aβ, microglia were responsible for fAβ clearance (Bacskai et al., 2001; Schenk et al., 1999). Transplantation of microglia into the hippocampus of Λβ42-injected Wistar rats also increased Aβ clearance (Takata et al., 2007).

Macrophage colony stimulating factor (MCSF) plays a critical role in the activation, maintenance and differentiation of microglia, the only cells in the brain that express the CSF receptor (CSFR1) (Erblich et al., 2011; Soulet and Rivest, 2008). Treatment of primary murine microglia with MCSF decreased their lysosomal pH, and increased fAβ degradation (Majumdar et al., 2007). This was attributable to increased lysosomal delivery of ClC-7, a chloride transporter necessary for complete lysosomal acidification (Majumdar et al., 2011). Consistent with this, MCSF treatment of AD mice (APP_Swe_/PS1) increased plaque clearance and improved performance in a spatial learning task (Boissonneault et al., 2009).

While some genetic risk factors for AD are known, causes of the late-onset form (LOAD) are still not understood (Kunkle et al., 2017; Lanoiselée et al., 2017; Tanzi, 2012). Clearance, rather than overproduction, of Aβ may be important for LOAD (Mawuenyega et al., 2010). Many enzymes have been described that degrade either monomeric or fibrillar Aβ. Addition of excess lysosomal enzymes to microglia led to increased degradation of fAβ (Majumdar et al., 2008). The lysosomal acidification caused by MCSF treatment (Majumdar et al., 2007) would also increase the activity of acid hydrolases.

Tripeptidyl peptidase 1 (TPP1), is a lysosomal enzyme candidate with a role in fAβ degradation (Dimitrova et al., 2017; Solé-Domènech et al., 2018). TPP1 is able to proteolyze fAβ efficiently at low pH (Solé-Domènech et al., 2018). Mass spectrometry analysis of peptides released from fAβ digested with TPP1 revealed several endoproteolytic cleavages including some within β-sheet regions that are important for fibril stability. Deficiency of TPP1 results in late infantile neuronal ceroid lipofuscinosis (LINCL), a childhood neurodegenerative lysosomal storage disease (Sleat et al., 1997). This enzyme is synthesized as an inactive zymogen that is stable at neutral pH. After incubation at low pH, the proenzyme rapidly undergoes autocatalytic activation (Golabek et al., 2008; Golabek et al., 2004; Guhaniyogi et al., 2009; Lin et al., 2001). The mature enzyme has a potent exopeptidase activity at pH 4 to 5, releasing tripeptides from proteins and oligopeptides with unsubstituted N-termini (Tian et al., 2006; Vines and Warburton, 1998). TPP1 also has endoproteolytic activity with a pH optimum of 3.0 (Ezaki et al., 2000).

In this study, we show that increasing TPP1 levels in primary microglia results in enhanced fAβ degradation. Conversely, in cells that normally degrade fAβ efficiently, reduction of TPP1 activity decreases fAβ degradation. Moreover, in a transgenic mouse model of AD we show that reduction of TPP1 activity leads to increased plaque deposition in both the cortex and hippocampus. These results highlight the potential importance of TPP1 in AD.

## 2. Materials and Methods

### 2.1 Materials

Chemical reagents were purchased from the following companies: Alexa 488-wheat germ agglutinin (A488-WGA), Invitrogen (Carlsbad, CA); Aprotinin, Sigma-Aldrich (St. Louis, MO); Λβ1-42 peptide (catalog # H 1368), Bachem (Torrance, CA); Avidin-biotin complex horseradish peroxidase (ABC-HRP) solution (Vectastain Elite ABC kit), Vector Laboratories (Burlingame, CA); Cy3 dye, Amersham, GE Healthcare Bio-Sciences (Piscataway, NJ); Deoxyribonuclease I (DNase I), Worthington Biochemical Corporation (Lakewood, NJ); Dextran conjugated to fluorescein and rhodamine (70,000 Dalton molecular weight), Molecular Probes, Invitrogen (Carlsbad, CA); Diaminobenzidine, Vector Laboratories; Leupeptin, Sigma-Aldrich; MCSF, R & D systems (Minneapolis MN); Pefabloc SC, Roche (Nutley, NJ); Pepstatin, Sigma-Aldrich; Shandon M1 solution, ThermoScientific (Waltham, MA); Trypsin (lyophilized powder), Worthington Biochemical Corporation. All other standard chemicals were purchased from Sigma-Aldrich unless otherwise stated. Purified recombinant human TPP1 was generated as described (Lin and Lobel, 2001). The tripeptidyl peptidase inhibitor Alanine-Alanine-Phenylalanine-Chloromethylketone, AAF-CMK, was from New England Biolabs (Ipswich MA).

### 2.2 Media

Dulbecco’s Modified Eagle Medium (DMEM), fetal bovine serum (FBS), L-glutamine, G418, McCoy’s 5A medium, and penicillin-streptomycin solution were purchased from Gibco Laboratories (Grand Island, NY).

### 2.3 RNAi Reagents

siRNA sequences were Dharmacon SMART siGenome on-target plus set of 4 duplexes and scrambled siRNA negative control from Thermo Scientific (Lafayette, CO). *Tpp1* targeting siRNAs were (1-GGGCUGAGUUUCAUCACUAUU, 2-CCUCUUCGGUGGCAACUUUUU, 3-ACUCAGACCUGGCUCAGUUUU, 4-GCACAUCAGGCAUCAGUAGUU). HiPerFect reagent was from Qiagen (Valencia CA).

### 2.4 Software

MetaMorph image analysis software was from MDS Analytical Technologies, Universal Imaging (West Chester, PA). Bar graphs and plots were made in GraphPad Prism version 8 software. P-values for the cell experiments were calculated using the Mann-Whitney U Test (GraphPad Prism). Actual P-values are reported unless they were below the maximum significant digits reported by the program, these are reported as P<1×10^−10^. P-values for the stained brain slices were calculated using GraphPad Prism Linear Regression ANCOVA method. The P value reported for the Thio-S stained brain slices is the difference in the slopes of the experimental lines. FIJI (Image J version 2.0.0-rc-64/1.51s) open source image processing software was used for processing of Thio-S brain slice images. Schematic designed with Inkscape version 0.48.

### 2.5 Cells

All cells were maintained at 37°C in a humidified 5% CO_2_ atmosphere. Primary mouse microglial cultures were isolated from the brains of one day old C57BL/6 mouse pups as described previously (Paresce et al., 1997). Briefly, brains were removed, minced with a scalpel blade in PBS, and sequentially digested with trypsin and DNase I. The resulting mixed glial culture was suspended in complete growth medium (DMEM, 10% FBS, 1% penicillinstreptomycin and 4mM L-glutamine) in a 75 cm^2^ tissue culture flask, cultured for two weeks and microglia isolated using the shaking method (Giulian and Baker, 1986; Nakajima, 1989). Microglia were plated at 70% confluency in complete growth medium on 35 mm coverslip-bottom poly-D-lysine coated dishes and cultured for 1-2 days prior to use for experiments. For the MCSF-treated microglia, the complete growth medium was supplemented with 25 ng/mL MCSF. MCSF medium was used for each medium exchange. After plating, cells were maintained in MCSF medium for the duration of the experiment.

J774.A1 mouse macrophage-like cells and human osteosarcoma U2OS cells were from American Type Culture Collection (Manassas, VA). J774 cells were maintained as adherent cultures in complete growth medium (see above) in 10 cm petri dishes (polystyrene, not treated for tissue culture). U2OS-SRA cells (Sanchez-Carbayo et al., 2003) stably express the murine scavenger receptor A (Pipalia et al., 2017). The expression of SRA by these cells leads to very strong attachment by the cells to tissue culture optimized plastic surfaces, so these cells were maintained in 10 cm (non-treated) petri dishes with McCoy’s 5A medium with 1.2 g/L sodium bicarbonate, 10% FBS, 1% penicillin/streptomycin. Selection pressure for SRA was maintained with 1 mg/mL G418.

### 2.6 fAβ Degradation Assay

Fluorescence microscopy images were collected on a Leica epifluorescence microscope with an oil immersion 40X NA 1.25 objective (Leica Microsystems, Wetzlar Germany) equipped with an Andor iXon cooled CCD camera. The microscope was controlled by MetaMorph Imaging system software (Molecular Devices Universal Imaging, West Chester, PA).

To track fAβ degradation, cells were incubated with fluorescently-labeled fibrillar Aβ42 (Chung et al., 1999). The fluorescence intensity in the cells was a measure of how much labeled fAβ remained in each set of cells. The loss of Cy3 signal indicated breakdown of the fAβ since Cy3 is quickly released from the cells if it is not attached to a larger protein (Paresce et al., 1997). This fluorescence degradation assay was comparable to degradation of radiolabeled fAβ (^125^I-fAβ) by microglia (Majumdar et al., 2008).

The Aβ1-42 peptide was derivatized with Cy3 mono-reactive dye according to the manufacturer’s instructions as described (Majumdar et al., 2007). Cy3-Aβ (7 μg/mL) was incubated in DMEM with 10 mg/mL bovine serum at 37°C for 1 hour. Negative stain electron microscopy at high magnification showed that a 1 μg/mL Cy3-Aβ solution formed fibrils with average diameters of 10 nm (Paresce et al., 1996). These Cy3-labeled fibrillar aggregates of Aβ are hereafter referred to as Cy3-fAβ. The Cy3-fAβ solution was incubated for 1 hour with cells plated on a 35 mm coverslip-bottom dish at 37°C. After 1 hour, excess Cy3-fAβ solution was washed off. Initial uptake by the cells was established for each experiment in three dishes fixed with 1% paraformaldehyde (PFA) immediately after 1 hour of loading with Cy3-fAβ (Majumdar et al., 2007). Experimental dishes were washed with complete medium and chased for various times with or without tested reagents. Cells were then fixed with 1% PFA, and the cell membranes were stained by incubation with 1 mg/mL A488-WGA in PBS for 10 minutes at room temperature. Cells were imaged by epifluorescence microscopy. Briefly, fields of cells were selected using the fluorescence from the A488-WGA stained cell borders. After selection, Cy3 images were acquired for each of these fields. Digital fluorescence images were analyzed using MetaMorph software. Images were background corrected, and the Cy3 integrated intensity measurement for each cell was recorded.

### 2.7 RNAi in U2OS-SRA cells

Scrambled (non-targeting) and *Tpp1* targeting siRNA sequences were purchased from Dharmacon as SMART siGenome on-target plus set of 4 duplexes. Stock solutions of these siRNAs (100 μM in RNase-free, pH 7.4 water) were prepared and frozen. U2OS-SRA cells were plated in 96 well plates at an initial density of 1 x 10^4^ cells/well. On the following day the cells were transfected according to Dharmacon’s standard transfection protocol. Individual or pooled 50 nM siRNAs complexed with HiPerFect reagent (Qiagen) were added to the plated cells for 72 hours in antibiotic free complete medium at 37°C in 5% CO_2_. After the transfection period, the medium in each well of the plate was changed and the cells assayed for ability to degrade and Cy3-fAβ as described above.

### 2.8 TPP1 Activity Assay

An endpoint TPP1 activity assay was conducted essentially as described (Sohar et al., 2000). Cell lysate (1.5 μg total protein) was incubated in a 100 μL substrate solution of 100 mM sodium acetate 150 mM sodium chloride, pH 4.5 buffer with 200 μM Ala-Ala-Phe 7-amido-4-methylcoumarin (AAF-AMC, Sigma), for 3 hours at 37°C. Each sample was split into 4 wells of a 384 well clear bottom polystyrene plate and read on a Spectramax M2 plate reader (Molecular Devices) at 37°C. The plate was read from the bottom using 351 nm excitation and 450 nm emission filters. The enzyme activity for each group of cells was expressed as the normalized value of arbitrary fluorescence units. All groups were normalized to the average fluorescence reading of the untreated cells.

### 2.9 Animals

All experiments were performed in compliance with the guidelines of the Institutional Animal Care and Use Committee of Robert Wood Johnson Medical School and Weill Cornell Medical College in accordance with the National Institutes of Health guidelines. All mice were in a C57BL/6 background. The J20 AD mouse model, B6.Cg-Tg(PDGFB-APPSwInd)20Lms/2J (Mucke et al., 2000), which expresses human APP with the Swedish (K670N/M671L) and Indiana (V717F) mutations (*APPSwInd*) under the control of the human *PDGFB* promoter, was acquired from Jackson Laboratories. Mice hemizygous for the J20 transgene are designated as “AD” or Tg+. These were crossed with C57BL/6 congenic *Tpp1* mutant mice (Sleat et al., 2008; Sleat et al., 2004). Mice homozygous for the neo^del^Arg446His *Tpp1* allele (designated *Tpp1*^f/f^) have <10 % TPP1 activity, synthesizing normal levels of properly spliced mRNA that encodes a missense mutation (Arg446His). Mice homozygous for the neo^ins^Arg446His *Tpp1* allele (designated *Tpp1*^-/-^) exhibit aberrant mRNA splicing and have <0.2 % TPP1 activity. In the absence of the J20 transgene (Tg-), the phenotype of all *Tpp1* genotypes (*Tpp1*^+/+, +/-, +/f and f/f^) exhibited no apparent differences up to the maximum age of 18 months (Sleat et al., 2008). Mice designated as having >50% TPP1 activity have at least one wild type *Tpp1* allele. Genotypes, TPP1 and Beta-galactosidase activities (normalized to protein levels, expressed as arbitrary units) were verified for all animals used in this study (J20, JAX protocol; TPP1and Betagalactosidase, Supplemental Table 1 (Sleat et al., 2008; Sleat et al., 2004)). Male and female mice were used for all experiments described (Supplemental Table 1).

### 2.10 Plaque Deposition in Mice

The following procedure was adapted from (Li et al., 2004). Tissue from the cerebrum was postfixed overnight in 4% PFA. The following day, the fixed brain was moved to a 30% sucrose solution at 4°C for at least 24 hours until the brain sunk to the bottom of the vial. The brains were snap frozen in Shandon M1 solution (ThermoScientific, Waltham MA), stored at −80°C, and maintained at −20°C overnight prior to sectioning. Sequential 40 μm coronal sections were cut through the frozen sections with a cryostat (Bright Instruments model OTF5000, Huntington, England). Slices were stored in 0.1M PB, 30% Ethylene Glycol, 30% sucrose, at −20°C in 24 well plates.

For fluorescence analysis of the plaques, slices from mouse brains 10 months (300 days) of age or older were stained with Thioflavine S (Thio-S, Sigma). Slices were washed with 0.1 M PB three times, stained in 0.005% Thio-S in 0.1 M PB for 15 minutes, then washed three times for 5 minutes each in 0.1M PB. Slices were mounted on glass slides and imaged with a Hamamatsu Nanozoomer at 20X at the NYULMC DART Experimental Pathology Research Lab, NYU Langone Medical Center.

Images of individual brain slices were opened and processed with FIJI software (Schindelin et al., 2012). A threshold was set to separate plaques from the background of the image. Plaques were scored based on minimum size criteria (22 μm) and circularity (0.2-1.0, calculated as: circularity = 4π(area/perimeter^2^), a value of 1.0 indicates a perfect circle). The region containing the hippocampus and cortex for each brain slice was hand drawn for each brain slice. The percent plaque coverage was determined based on the area covered by plaques in the region divided by the defined area of the cortex or hippocampus (Measure function in FIJI). The plaque count was determined using the Analyze Particles function of FIJI. All numbers were exported to Excel, separated by individual slice.

## 3. Results

### 3.1 Uptake of TPP1 by microglia enhances degradation of fAβ

Exogenously added recombinant TPP1 is internalized by cells (via the mannose-6-phosphate receptor) with an apparent EC_50_ for uptake of 1.5 nM (Lin and Lobel, 2001). To determine if increased levels of TPP1 would enhance degradation of fAβ, we loaded mouse microglia with Cy3-fAβ for one hour and then incubated the cells with or without 100 nM recombinant TPP1 for 72 hours. Untreated microglia degraded negligible amounts of the Cy3-fAβ during this time, but the additional TPP1 led to degradation of more than half of the internalized Cy3-fAβ (Figure 1).

**Figure 1.**
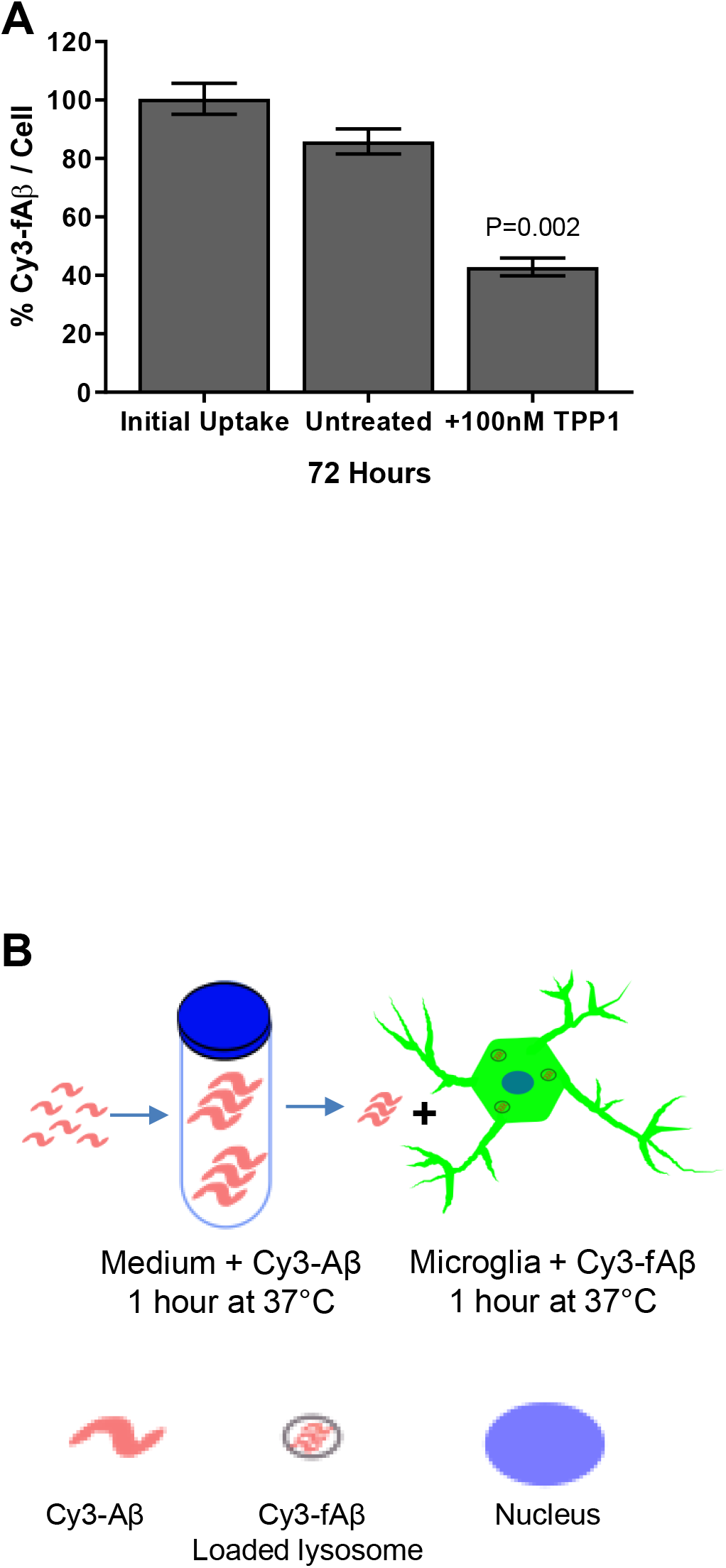
Addition of TPP1 to microglia enhances fAβ degradation. Primary murine microglia were incubated with Cy3-fAβ for one hour and, where indicated, treated with 100 nM purified proTPP1 for the duration of a 72 hour chase. Cells were PFA fixed and imaged by epifluorescence microscopy. The Cy3-fAβ intensity was measured for each cell for ~190 total cells (total from 3 separate experiments). Within 72 hours, treated microglia efficiently degraded fAβ, as measured by loss of Cy3 fluorescence, while degradation by untreated cells was limited. Bars are mean ± SEM for all cells measured (P=0.002). (B) Schematic of loading Cy3-fAβ to microglia.

### 3.2 Tripeptidyl peptidase activity is required for efficient degradation of fAβ

Macrophages and MCSF-activated microglia can efficiently degrade fAβ (figure 2). We tested whether tripeptidyl peptidase inhibition would impact fAβ degradation by these cells. We also tested this in an osteosarcoma cell line that stably expressed Scavenger Receptor A (U2OS-SRA) (Pipalia et al., 2017; Sanchez-Carbayo et al., 2003). The inhibitor Alanine-Alanine-Phenylalanine-Chloromethylketone (AAF-CMK) inhibits TPP1 (Junaid et al., 2000; Vines and Warburton, 1998) and TPP2 (Lévy et al., 2002). Fifty micromolar AAF-CMK was added to the growth medium of all three cell types for 1 hour before Cy3-fAβ was added to the cells. The cells were then chased in medium with or without the inhibitor for 48 hours after Cy3-fAβ uptake. This led to a nearly complete inhibition of Cy3-fAβ degradation as measured by release of Cy3 from the cells (Figure 2A-C). Untreated cells degraded 60-75% of internalized Cy3-fAβ within 48 hours.

**Figure 2.**
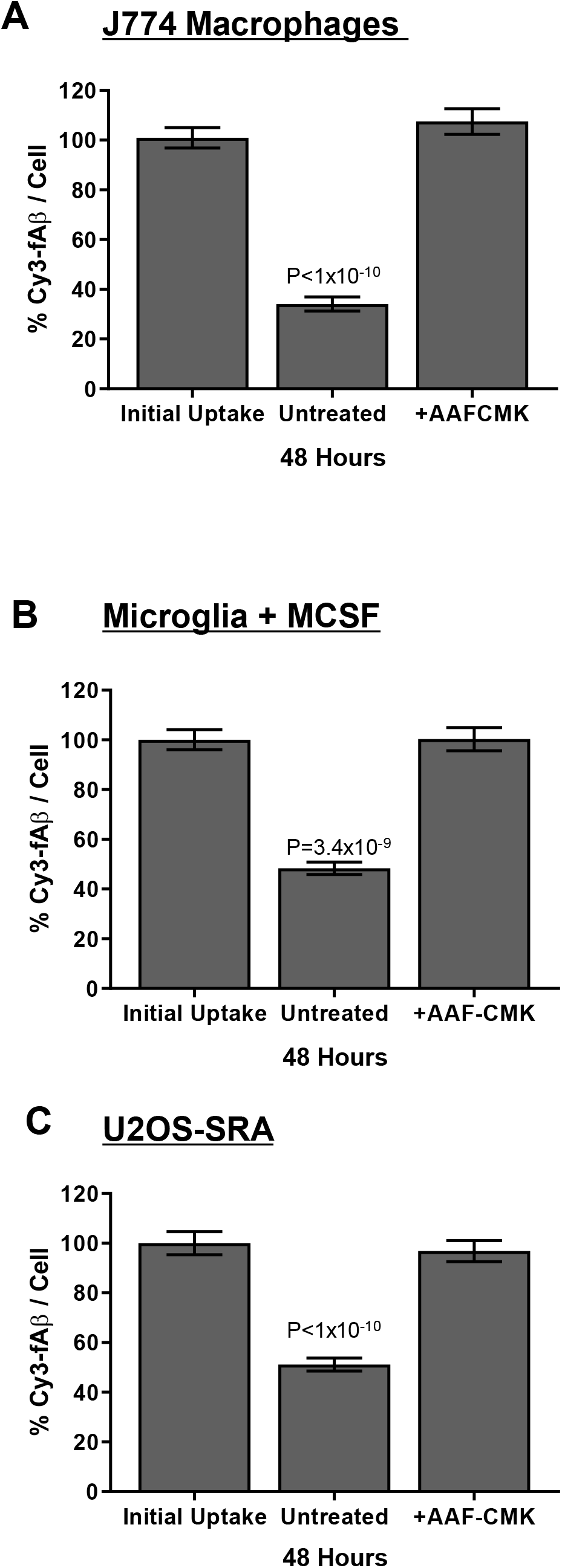
Degradation of internalized fAβ is blocked by a tripeptidyl peptidase inhibitor. J774A.1 macrophage cells (A) (P<1×10), microglia treated with MCSF (B) (P=3.4×10^−9^) and U2OS-SRA cells (C) (P<1×10^−10^) were pre-treated for 1 hour with or without 50 μM AAF-CMK. Subsequently, the cells were loaded with Cy3-fAβ for one hour and chased for 72 hours with or without AAF-CMK in the medium. Cells were PFA fixed and imaged by epifluorescence microscopy. The Cy3-fAβ intensity was measured for each cell for ~120 total cells (total from 3 replicate experiments) per condition. The inhibitor-treated cells showed reduced fAβ degradation. Bars are mean ± SEM for all cells measured.

### 3.3 RNAi knockdown of TPP1 reduces fAβ degradation

We further tested the role of TPP1 in fAβ degradation by reducing its expression in cells by RNA interference (RNAi). In these experiments, we used U2OS-SRA cells due to the difficulty in efficiently transfecting J774 macrophage cells or microglia. The U2OS-SRA cells degrade Cy3-fAβ almost as well as J774 macrophages or MCSF-treated microglia (Figure 2). The SRA expression by this cell line allows it to take up fAβ and deliver it to lysosomes (Majumdar et al., 2011).

We tested multiple siRNA sequences that targeted *Tpp1* and pooled two to knock down TPP1 expression in these cells. The TPP1 activity in cells treated with these siRNAs for 72 hours was about 30% of the activity in untreated cells or cells treated with scrambled siRNA (Figure 3A). The cells were incubated for 72 hours with the siRNAs (or controls) and then loaded with Cy3-fAβ for one hour and chased for 24 hours. The initial uptake of Cy3-fAβ by untreated and siRNA treated cells was pooled since the different conditions did not alter uptake. After siRNA treatment, Cy3-fAβ degradation was inhibited almost completely compared to control U2OS-SRA cells (Figure 3B). Two sets of controls were used in these experiments. One set of cells was treated with non-specific (scrambled) siRNA. The other set was not treated with any siRNA. Both the untreated cells and those treated with scrambled siRNA degraded more than half of the Cy3-fAβ after a 24 hour chase period. Thus, loss of TPP1 expression greatly reduces degradation of internalized fAβ.

**Figure 3.**
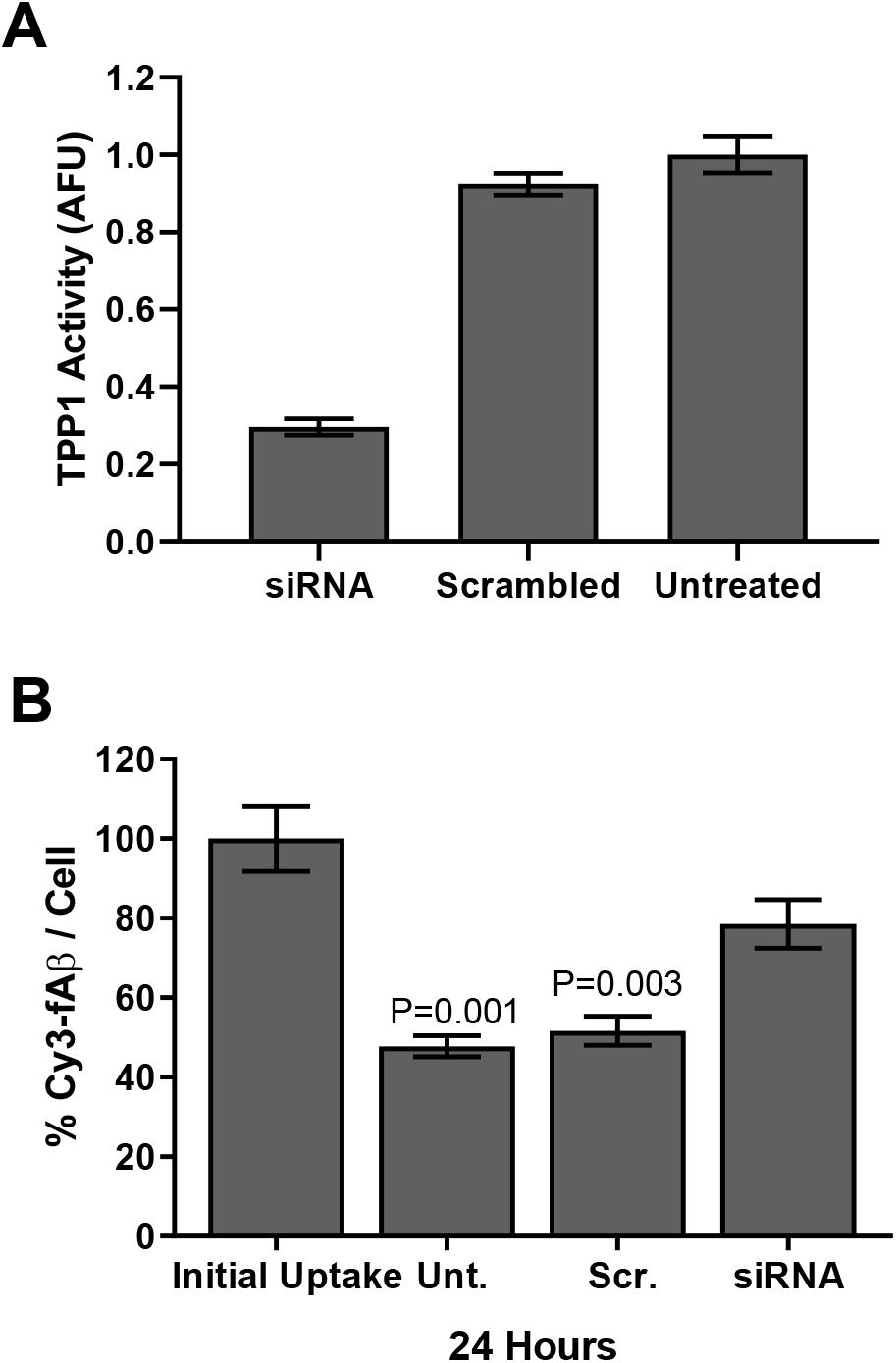
Degradation of fAβ is blocked by RNAi knockdown of *Tpp1* in U2OS-SRA cells.. (A) TPP1 activity in cells incubated for 72 hours with indicated siRNA was measured in cell lysates. Enzyme activity (shown as arbitrary fluorescence units, AFU) was normalized to untreated cells (P=0.0002). (B) The Cy3-fAβ intensity was measured in untreated, mock (scrambled) siRNA-treated, and *Tpp1* targeted siRNA-treated cells incubated with Cy3-fAβ followed by a 24 hour chase (P=0.0008). Uptake was not altered by RNAi treatment, so the combined average initial uptake is shown. Only the cells treated to knock down TPP1 expression did not degrade fAβ efficiently. Approximately 500 fixed cells (total for all conditions) were imaged by epifluorescence microscopy. Bars are mean ± SEM for all cells measured.

### 3.4 Microglia from *Tpp1^-/-^* mice show impaired degradation of fAβ

Previous studies indicate that MCSF treatment of wild type microglia significantly enhances their ability to degrade fAβ (Majumdar et al., 2007). MCSF treated microglia have more acidic lysosomes (Majumdar et al., 2007). This lower lysosomal pH creates a more optimal environment for the function of lysosomal enzymes. To investigate the participation of TPP1 in this process, we performed parallel experiments on primary microglia from wild type and *Tpp1*^-/-^ mice. The TPP1-deficient microglia were only able to degrade ~15% of the endocytosed fAβ in 72 hours as compared to ~70% degradation in the wild type microglia (Figure 4).

**Figure 4.**
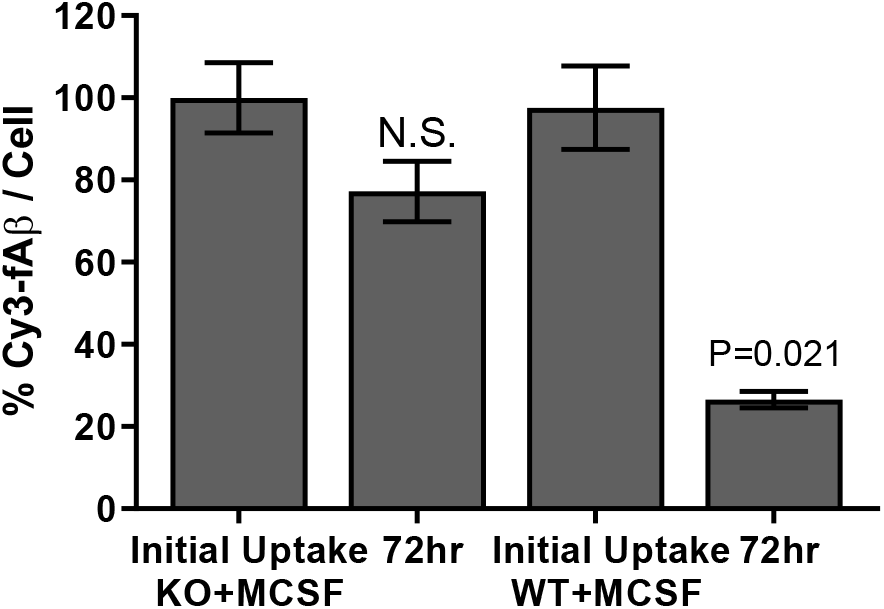
MCSF-treated *Tpp1*^-/-^ microglia do not degrade fAβ efficiently.. Microglia (MG) isolated from wildtype or *Tpp1*^-/-^ mice pretreated with MCSF for 2 weeks were incubated with Cy3-fAβ (1 hr) followed by a 72 hr chase in MCSF-containing medium. The Cy3-fAβ intensity was measured by epifluorescence microscopy for each fixed cell (50 cells per experiment, 3 replicates from one MG prep, for total ~150 cells per condition). MCSF-treated wild type microglia efficiently degraded fAβ (P=0.021) compared to limited fAβ degradation by MCSF-treated *Tpp1*^-/-^ microglia. Bars are mean ± SEM for all cells measured.

### 3.5 Effect of TPP1 on Plaque Deposition in an AD Mouse Model

We crossed J20 AD model mice (designated Tg+) with strain-matched lines that had reduced TPP1 expression levels. If TPP1 expression is important for the degradation of Aβ plaques, mice that lack normal levels of expression should have increased Aβ plaque load. *Tpp1*^f/f^ mice have <10% of wild-type TPP1 activity (Supplemental Table 1). The two groups of control animals were Tg+ with 50% of normal expression of TPP1 (Tpp1 ^+/f^) and mice that did not express the APP transgene (Tg-). For these non-AD mice, *Tpp1*^+/+^, *Tpp1*^+/f^, and *Tpp1*^f/f^ animals did not have plaque deposition and thus we pooled mice with all levels of TPP1 expression for the non-AD group.

We labeled sections with Thio-S to stain the amyloid plaques (Figure 5A). Quantitative comparisons of the Thio-S labeled plaques in the brains of the mice aged 300 days or older showed increased plaque coverage of the cortex and hippocampus with age, and mice with limited TPP1 activity expression had greater plaque load than mice with TPP1 activity of 50% or more (Figure 5, Supplemental Table 1). The percent of both the cortex and hippocampus covered by plaques was significantly higher in mice with limited TPP1 activity compared to mice that had TPP1 activity of 50% or more (Figure 5B and D). The number of plaques increased with age in both the cortex and hippocampus (Figure 5C and E). Plaque numbers were significantly greater in the cortex and hippocampus of mice with limited TPP1 activity.

**Figure 5.**
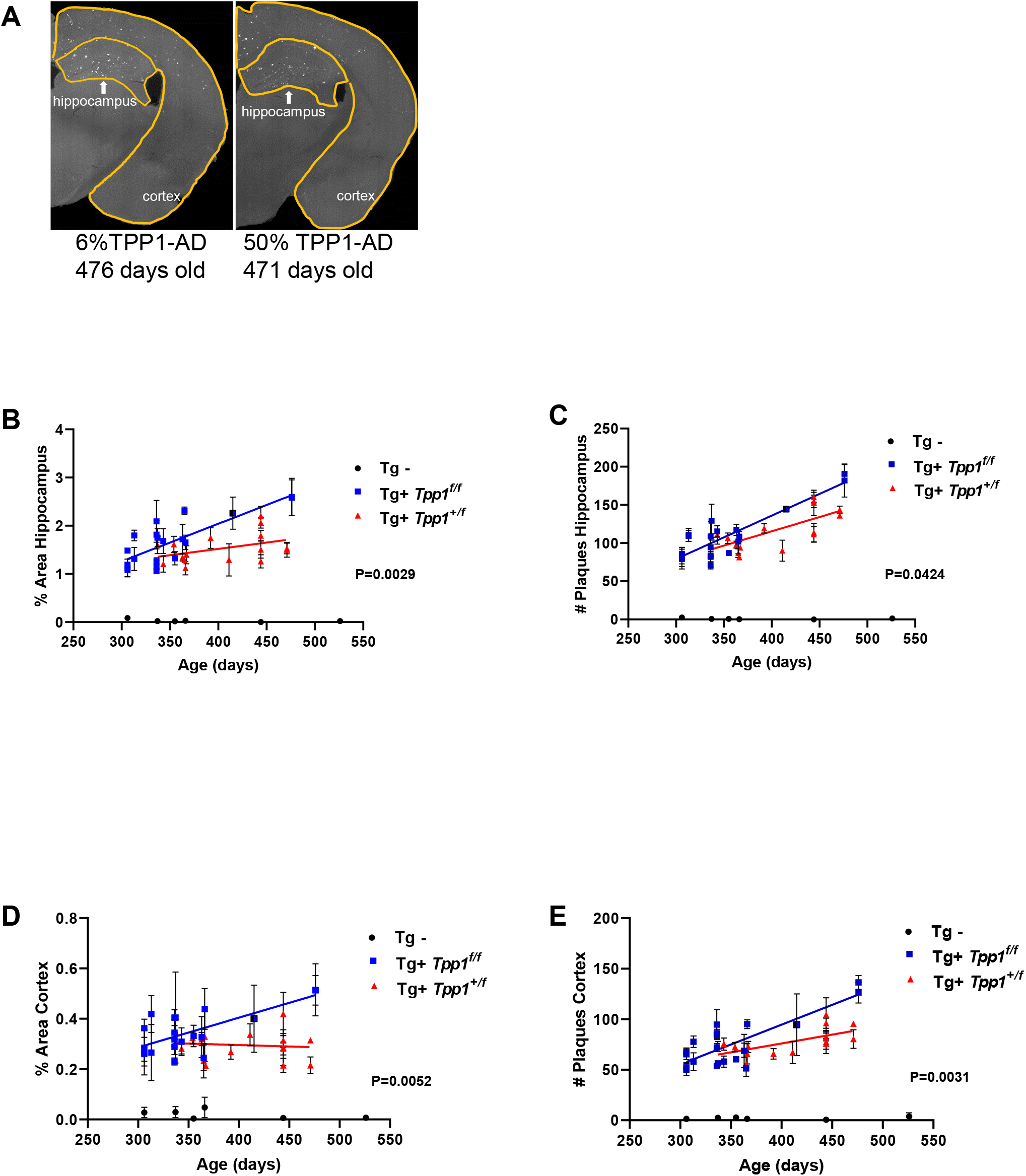
Reduced TPP1 activity increases plaque burden in AD mice.. (A) Amyloid plaques in the brain were visualized by Thio-S staining in Tg+ and Tg-mice that had <10% or 50+% TPP1 activity. Representative images of older Thio-S stained brain slices. (B) Percent plaque coverage of the hippocampus in mice age 300 days and older. Square blue symbols are Tg+ animals with <10% TPP1 activity. Red triangles are Tg+ mice with 50% TPP1 activity. Bars show the SEM of the values from 3 individual slices (solid symbols, split symbols indicate animals for which only 2 slices were analyzed), P=0.0029. (C) The total number of Thio-S labeled plaques from the hippocampus of brain slices of mice 300 days of age and older. Symbols and bars are as described above (P=0.0424). (D) Percent plaque coverage of the cortex in mice age 300 days and older. Symbols and bars are as described above (P=0.0052). (E) The total number of Thio-S labeled plaques from the cortex of brain slices of mice 300 days of age and older. Symbols and bars are as described above (P=0.0031). Statistical analysis conducted in GraphPad Prism 8.0 comparing Tg+ groups (ANCOVA Linear Regression), blue line for Tg+ animals with <10% TPP1 activity and red line for the Tg+ mice with 50% TPP1 activity.

## 4. Discussion

In Alzheimer’s disease, the net accumulation of Aβ is due to an imbalance between its production and degradation. Several mutations in humans and animal models increase the rate of Aβ production (Saido and Leissring, 2012). Most of the early-onset autosomal dominant AD results from pathogenic genetic mutations in APP that alter its proteolytic processing and endolysosomal transport, leading to increased Aβ production (Kunkle et al., 2017; Lanoiselée et al., 2017). Other mutations that result in increased production of Aβ and that have been associated with increased incidence of AD include PSENs 1 and 2, components of the γ-secretase complex, and BACE 1. These mutations increase plaque burden and are associated with increased risk of dementia.

The regulation of Aβ degradation is still an active area of research. The Aβ peptide is produced mostly by neurons and is found both inside cellular organelles and in the extracellular space (Dominguez et al., 2005). Aβ spontaneously forms β-sheet fibrils that can aggregate and associate with other proteins to form plaques (Glenner and Wong, 1984). It is then taken up by many cell types and accumulates in their lysosomes.

There are several proteolytic enzymes that have been implicated in degradation of Aβ (reviewed in (Saido and Leissring, 2012)). Decreasing the levels of several of these in mouse models resulted in increased endogenous cerebral Aβ, which was additive and gene dose dependent. Thus, more than one enzyme may be critically required for Aβ degradation, and there may be little reserve capacity for the clearance of Aβ in the brain (Saido and Leissring, 2012).

The β-sheet structure of fibrillar Aβ makes it protease resistant, protecting most of the sequence from degradative enzymes. Disruption of the β-sheet would provide access to the peptide bonds that are cleaved by lysosomal proteases. The most studied lysosomal enzymes in AD, Cathepsins B and D, were found in extracellular Aβ plaques (Cataldo and Nixon, 1990; Mueller-Steiner et al., 2006). Cathepsin B is able to cleave fibrillar Aβ with multiple cut sites just before and within the β-sheet region both *in vivo* and *in vitro* (Mueller-Steiner et al., 2006). Cathepsin B knock-out mice that also expressed human APP had increased hippocampal Aβ plaque load (Mueller-Steiner et al., 2006). Upregulation of cathepsin B in hippocampal neurons of APP/PS1 transgenic mice reduced Aβ plaque load and showed memory improvement (Embury et al., 2017). Cystatin C, a cysteine protease inhibitor, has also been genetically linked to late-onset AD (Bertram et al., 2007; Sun et al., 2008). In mice, downregulation of cystatins led to reduced Aβ1-42 levels (Sun et al., 2008; Wang et al., 2012; Yang et al., 2011). Cathepsin D, an aspartic protease, cuts Aβ42 within the β-sheet region (Hamazaki, 1996; Mackay et al., 1997).

The lysosomal protease TPP1 has endopeptidase activity in addition to its function in removing tripeptides from the unsubstituted N-termini of proteins and peptides. *In vitro* studies indicate that TPP1 cleaves fibrillar Aβ at numerous sites including those in the β-sheet region (Solé-Domènech et al., 2018). TPP1 and Cathepsin B have overlapping cut sites. Some of these cleavages are within the KLVFF hydrophobic core of the β-sheet and are the most likely to destabilize it. Further degradation of Aβ fibrils can then occur due to the action of many lysosomal enzymes after the protective β-sheet structure has been disrupted.

We found that delivery of excess TPP1 to microglial lysosomes by endocytosis of exogenous enzyme increased the ability of unactivated microglia to degrade fAβ. Even though the endoprotease activity of each TPP1 enzyme would remain low at the pH of microglial lysosomes, this can be compensated by a large increase in the abundance of the enzyme. Although unactivated microglia are very poor at degrading fAβ, several other cells types (including macrophages, activated microglia, and U2OS-SRA cells) can degrade fAβ in their lysosomes (Majumdar et al., 2008; Majumdar et al., 2007). Inhibition of TPP1 in these three cell types with the tripeptidyl peptidase inhibitor AAF-CMK led to almost complete loss of fAβ degradation. Similarly, siRNA reduction of TPP1 expression greatly reduced fAβ degradation in U2OS-SRA cells. Thus, levels of TPP1 activity play a critical role in the degradation of fAβ. It is not clear which role each enzyme plays, but it is possible that multiple cuts in the Aβ attached to a fibril are required to release it efficiently.

We present data in the J20 transgenic mouse model of AD that show TPP1 activity is also important for degradation of amyloid *in vivo*. There was a significant increase in the amyloid burden of Tg+ mice with <10% of normal TPP1 activity. (Mice with ≤3% TPP1 do not survive long enough to develop plaques.)

Currently, there are no reported associations of variations in TPP1 and AD in humans. However, as a member of the CLEAR (Coordinated lysosomal expression and regulation) network of genes regulated by TFEB (transcription factor EB), TPP1 expression increases with TFEB activation (Song et al., 2013), and modulation of lysosomal function by TFEB has been proposed as a potential target for treatment of AD based on animal results (Xiao et al., 2014). TPP1 levels vary with age and cell type. In the human hippocampus, the CA2/3 regions show the strongest immunoreactivity at a young age that weakens for adults but returns to higher levels at advanced age. The pyramidal neurons of the CA1 region also have weaker TPP1 expression than CA2/3 at all ages (Kida et al., 2001).

Based on multiple lines of evidence in this study that implicate TPP1 in the clearance of fAβ, elevating levels of this enzyme in microglia and/or other cells in the brain may be beneficial in reducing fAβ levels in the AD brain. This is particularly exciting given that cerebroventricular administration of TPP1 has been shown to be effective as an enzyme replacement therapy for LINCL (Schulz et al., 2018). However, the utility of this approach for AD remains to be demonstrated and it is possible that there are indirect effects arising from reduction of TPP1 in our *in vivo* studies. A transgenic mouse model has been developed that constitutively overexpresses TPP1 10-20 fold higher than wild-type levels (Nemtsova et al., 2018). This should allow direct evaluation of the potential therapeutic utility of TPP1 in fAβ clearance and amelioration of the AD phenotype.

## Acknowledgements

The authors thank the laboratory of Dr. Lennart Mucke of The J. David Gladstone Institutes of San Francisco, CA for the use of the PDGF-hAPP (J20) transgenic mice. The authors would also like to thank Drs. Branka Brukner Dabovic and Cynthia Loomis of the NYULMC DART Experimental Pathology Research Lab (NYU Langone Medical Center) for imaging services. We also thank Harold Ralph and Dr. Sushmita Mukherjee (Weill Cornell Medicine) for assistance with automated microscopy and image analysis.

---

**Supplemental Table 1.** Genotype, Age, Sex, And Enzyme Activity of mice used for Thio-S plaque evaluation in figure 5. Enzyme activity is normalized to protein levels and is expressed as arbitrary units.

| Genotype | Animal Identifier | Age (Days) | Sex | % Plaque (Avg) Hip | % Plaque (Avg) Cor | # Plaque (Avg) Hip | # Plaque (Avg) Cor | TPP1 | β-galactosidase |
| --- | --- | --- | --- | --- | --- | --- | --- | --- | --- |
| Tg+ (f/f) | 6736-37A | 306 | F | 1.08 | 0.29 | 79.00 | 79.00 | 7.4 | 307 |
| Tg+ (f/f) | 6734 | 306 | F | 1.49 | 0.26 | 83.67 | 83.67 | 6.9 | 298 |
| Tg+ (f/f) | 6729 | 306 | M | 1.13 | 0.27 | 86.00 | 86.00 | 7.5 | 276 |
| Tg+ (f/f) | 6733 | 306 | M | 1.19 | 0.36 | 81.67 | 81.67 | 7.1 | 307 |
| Tg+ (f/f) | 7109 | 313 | F | 1.31 | 0.27 | 109.33 | 109.33 | 7.4 | 293 |
| Tg+ (f/f) | 7102 | 313 | M | 1.80 | 0.42 | 110.67 | 110.67 | 8.2 | 300 |
| Tg+ (f/f) | 6720 | 336 | F | 1.15 | 0.32 | 82.00 | 82.00 | 8.4 | 327 |
| Tg+ (f/f) | 6723 | 336 | F | 2.09 | 0.35 | 70.67 | 70.67 | 7.8 | 318 |
| Tg+ (f/f) | 6724 | 336 | F | 1.07 | 0.29 | 94.33 | 94.33 | 7.2 | 305 |
| Tg+ (f/f) | 6717 | 336 | M | 1.27 | 0.23 | 83.33 | 83.33 | 8.7 | 327 |
| Tg+ (f/f) | 6718 | 336 | M | 1.82 | 0.40 | 108.67 | 108.67 | 8.7 | 306 |
| Tg+ (f/f) | 7977 | 337 | M | 1.75 | 0.41 | 129.33 | 129.33 | 8.6 | 334 |
| Tg+ (f/f) | 7969 | 343 | M | 1.68 | 0.31 | 115.33 | 115.33 | 9.1 | 335 |
| Tg+ (f/f) | 7856-58C | 355 | M | 1.32 | 0.33 | 87.00 | 87.00 | 9.1 | 352 |
| Tg+ (f/f) | 7755-57C | 363 | M | 1.72 | 0.33 | 117.33 | 117.33 | 10.0 | 379 |
| Tg+ (f/f) | 7846 | 365 | F | 2.31 | 0.25 | 102.00 | 102.00 | 9.1 | 366 |
| Tg+ (f/f) | 7745A | 366 | F | 1.66 | 0.44 | 108.33 | 108.33 | 65.3 | 330 |
| Tg+ (f/f) | 6444 | 415 | M | 2.26 | 0.40 | 144.50 | 144.50 | 8.4 | 337 |
| Tg+ (f/f) | 6002 | 476 | F | 2.60 | 0.51 | 190.67 | 190.67 | 9.0 | 348 |
| Tg+ (f/f) | 6005 | 476 | F | 2.58 | 0.51 | 182.00 | 182.00 | 8.4 | 328 |
| Tg+ (f/+) | 7976 | 337 | M | 1.60 | 0.35 | 108.00 | 108.00 | 66.6 | 327 |
| Tg+ (f/+) | 7968 | 343 | M | 1.20 | 0.28 | 110.67 | 110.67 | 62.7 | 300 |
| Tg+ (f/+) | 7947 | 354 | M | 1.61 | 0.32 | 106.33 | 106.33 | 68.7 | 340 |
| Tg+ (f/+) | 7755-57A | 363 | M | 1.35 | 0.32 | 98.00 | 98.00 | 74.4 | 351 |
| Tg+ (f/+) | 7840-7889 | 365 | F | 1.29 | 0.23 | 86.67 | 86.67 | 65.4 | 343 |
| Tg+ (f/+) | 7745B | 366 | F | 1.12 | 0.33 | 81.33 | 81.33 | 10.0 | 387 |
| Tg+ (f/+) | 5974 | 367 | M | 1.40 | 0.21 | 94.00 | 94.00 | 60.8 | 283 |
| Tg+ (f/+) | 6710 | 392 | F | 1.75 | 0.27 | 119.33 | 119.33 | 58.8 | 284 |
| Tg+ (f/+) | 6725 | 411 | M | 1.29 | 0.34 | 90.00 | 90.00 | 58.1 | 281 |
| Tg+ (f/+) | 5545-48A | 444 | F | 1.26 | 0.21 | 113.67 | 113.67 | 73.1 | 310 |
| Tg+ (f/+) | 5545-48B | 444 | F | 1.80 | 0.31 | 152.67 | 152.67 | 66.6 | 309 |
| Tg+ (f/+) | 5545-48D | 444 | F | 1.50 | 0.28 | 111.33 | 111.33 | 76.8 | 335 |
| Tg+ (f/+) | 5543-44A | 444 | M | 2.20 | 0.29 | 154.67 | 154.67 | 68.6 | 332 |
| Tg+ (f/+) | 5543-44B | 444 | M | 2.06 | 0.42 | 160.00 | 160.00 | 53.6 | 349 |
| Tg+ (f/+) | 5984-85A | 471 | F | 1.50 | 0.31 | 136.00 | 136.00 | 65.5 | 305 |
| Tg+ (f/+) | 5984-85B | 471 | F | 1.53 | 0.22 | 143.33 | 143.33 | 60.2 | 276 |
| Tg- (f/f) | 6736-37B | 306 | F | 0.08 | 0.03 | 2.67 | 2.67 | 7.6 | 306 |
| Tg- (f/+) | 7974 | 337 | M | 0.02 | 0.03 | 0.67 | 0.67 | 63.7 | 316 |
| Tg- (f/f) | 7856-58B | 355 | M | 0.02 | 0.00 | 0.67 | 0.67 | 9.4 | 356 |
| Tg- (f/f) | 7847-48A | 366 | M | 0.03 | 0.05 | 0.33 | 0.33 | 9.6 | 353 |
| Tg- (+/-) | 5545-48C | 444 | F | 0.00 | 0.01 | 0.00 | 0.00 | 65.7 | 278 |
| Tg- (+/+) | 5023-24-25 | 526 | F | 0.02 | 0.01 | 1.33 | 1.33 | 115.7 | 258 |

